# Accurate registration of 3D time-lapse microscopy images

**DOI:** 10.1101/436824

**Authors:** Seyed M. M. Kahaki, Shih-Luen Wang, Armen Stepanyants

## Abstract

*In vivo* imaging experiments often require automated detection and tracking of changes in the specimen. This problem, however, can be hindered by variations in the position and orientation of the specimen relative to the microscope, as well as by linear and nonlinear deformations. Here, we present a feature-based registration method, coupled with translation, rigid, affine, and B-spline transformations, designed to address these issues in 3D time-lapse microscopy images. In this method, features are detected as local intensity maxima in the source and target image stacks, and their similarity matrix is used as an input to the Hungarian algorithm to establish initial correspondences. Random Sampling Consensus algorithm is then employed to eliminate outliers. The resulting set of corresponding features is used to determine the optimal transformations. Accuracy of the proposed algorithm was tested on fluorescently labeled axons imaged over a 48- day period with a two-photon laser scanning microscope. For this, multiple axons in individual stacks of images were traced semi-manually in 3D, and the distances between the corresponding traces were measured before and after the registration. The results show that there is a progressive improvement in the registration accuracy with increasing complexity of the transformations. In particular, an accuracy of less than 1 voxel (0.26 μm) was achieved with a regularized B-spline transformation. To illustrate the utility of the proposed method, registered images were used to automatically track synaptic boutons on axons over the entire duration of the experiment, yielding 99% precision and recall.

## INTRODUCTION

Accurate registration is often required for the analysis and visualization of medical images. For instance, 3D optical microscopy imaging of a large region of interest (ROI) is often performed by acquiring stacks of images that tile the ROI with overlaps. The same ROI can, in addition, be imaged multiple times in a time-lapse manner. Because such imaging experiments are often done *in vivo*, registration is required to eliminate artifacts related to tissue translation, rotation, as well as linear and non-linear distortions. Three of the most common registration problems include (*i*) registration of image planes within individual stacks, (*ii*) registration of image stacks within a larger ROI, and (*iii*) registration of image stacks over time. There are numerous available registration algorithms [1], however, in our experience, they do not reliably yield sub-micrometer accuracy required in many biological experiments. In particular, structural changes in the brain, related to learning and memory formation, can be visualized in *in vivo* imaging experiments, but automated detection and analyses of these changes are hindered by the relatively low accuracy of the existing registration methods. Therefore, we propose a new method, capable of achieving a sub-micrometer accuracy. This method was validated on a dataset of two-photon laser-scanning microscopy images of mouse brain acquired *in vivo*, and the registration results were used to reliably track synaptic structures over the 48-day duration of the experiment.

## METHODS

To register two time-lapse stacks of images, referred to as source and target, we detect intensity-based features in the two stacks, match these features, and use the matches to determine the optimal registration transformation. Features in this study are defined as small volumes (9×9×9 voxels for the data in Figure 1) centered at local intensity maxima in the stacks. Other features, such as SIFT, SURF, and BRIEF [2], may be used instead. Prior to the detection of the maximum intensity voxels, the stacks are filtered with a Gaussian filter (1×1×1 voxels in size) to reduce the effect of noise. To have a uniform distribution of features, we use a sequential algorithm in which after the detection of one local maximum, voxels belonging to the corresponding feature are eliminated from consideration, and the algorithm continues until there are no new features left.

**Figure 1:**
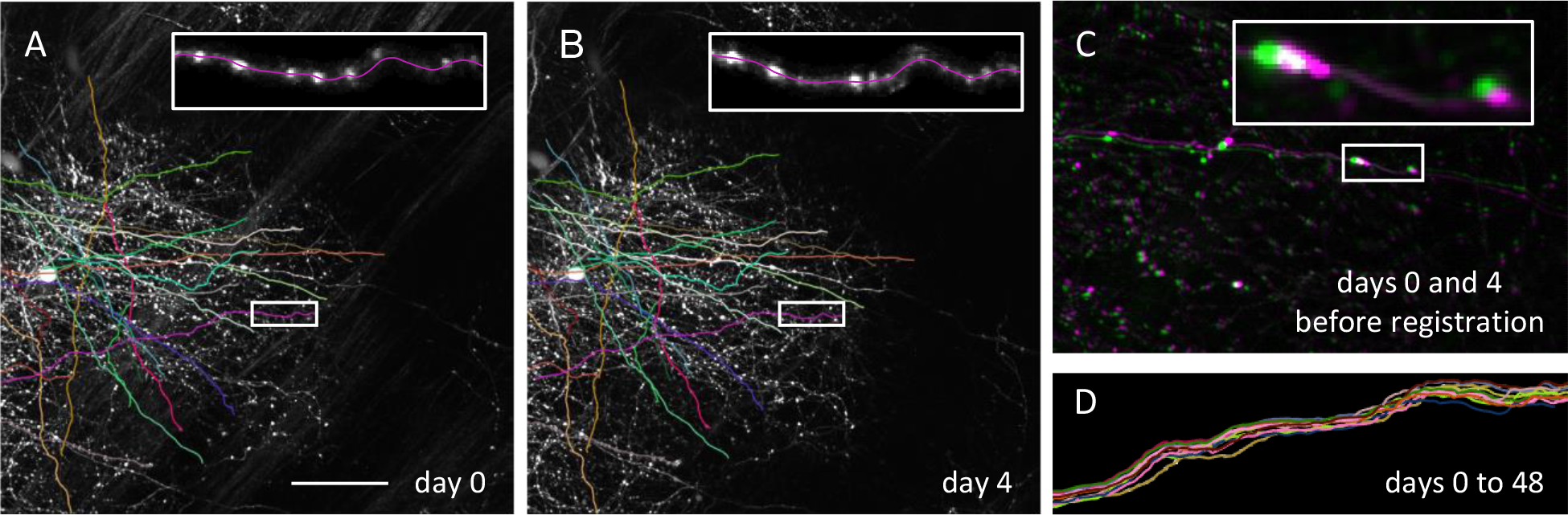
Registration is required for automated analyses of *in vivo* time-lapse images of neurons. **A, B.** Maximum intensity projections of two image stacks acquired with a 4-day interval, showing fluorescently labeled axons of cortical neurons. Traces of some of the axons obtained with NCTracer software are shown with colored lines. Insets in A and B show zoomed-in views of an axon with clearly visible structural changes in axonal boutons (bright spots).**C.** Overlap of A and B based on the alignment achieved during the experiment illustrates significant displacement and distortion of the tissue between imaging sessions (red-green). **D.** Overlap of 13 traces of an axon imaged with a 4-day interval shows significant misalignment. Scale bar in (A) corresponds to 50 μm for (A) and (B), and 25 μm for (C) and (D).

To establish initial matches between the features in the source and target stacks, we compute a matrix of feature similarities, *S*, in which an element *s*_*ij*_ denotes the correlation coefficient between the feature *i* in the source stack and the feature *j* in the target stack. Matrix *S* is used as an input to the Hungarian algorithm [3] to determine the initial correspondences between the features. In practice, the initial matches of features often contain outliers, and if not removed, even a small fraction of outliers can significantly reduce the registration accuracy. Therefore, we developed a method to eliminate outliers by combining the Random Sample Consensus (RANSAC) algorithm [4] with a specific class of optimal 3D transformations. In this procedure, a subset of corresponding features is randomly selected, and an optimal transformation is calculated to register this subset. One pair of features is used for translation, three for rigid, four for affine, and six for B-spline transformation. The detected optimal transformation is then applied to all source features, and those that ended up within 3 voxels from their corresponding target pairs are deemed to be inliers. This process continued until the maximum number of sampling steps is reached, and the largest set of inlier matches is used for the registration.

The optimal registration transformation, *f*, is determined by minimizing the mean-square-distance between the positions of the transformed source feature centers, *X*_*i*_, and the corresponding target feature centers, *Y*_*i*_. Below we consider and compare the following four classes of transformations: translation, rigid, affine, and B-spline [5]. The latter two are regularized by the elastic distortion energies associated with the transformations:

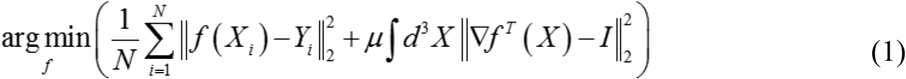

In this expression, *N* is the number of corresponding inlier features, *I* denotes the 3×3 identity matrix, and *μ* is referred to as the regularization strength. Eq. (1) is solved analytically to find the optimal translation, rigid, and affine transformations, and numerically with the gradient descent method in the case of B-spline.

## RESULTS

To evaluate the accuracy of the registration method we applied it to *in vivo* time-lapse images of fluorescently labeled axons in mouse barrel cortex obtained as part of another study [6]. The image stacks, 270×270×250 μm^3^, were acquired at a voxel volume of 0.26×0.26×0.80 μm^3^ with a 4-day interval in thirteen imaging sessions. A subset of axons contained in the images was traced and optimized with NCTracer software [7,8]. Boutons on these axons were detected and tracked over time with BoutonAnalyzer software [6]. Figures 1A, B show maximum-intensity projections of two image stacks of the same brain region acquired on days zero and four. A custom-built system was used to align the animal’s head during imaging, but residual misalignment is clearly visible in the image overlap (Figure 1C) and in the overlap of axonal traces (Figure 1D).

### Validation of the registration method

Registration can significantly improve on the original alignment (Figure 2A, B). To evaluate the registration accuracy, the average distances between the corresponding traces in different time-lapse images were calculated before and after registration (Figure 2). The results show that significant improvement over the original alignment (12.3 voxels, Figure 2D) is achieved with all four transformation types. The average trace distance decreased gradually with increasing complexity of the transformations, and the highest accuracy (0.7 voxels or 0.2 μm) was attained with the regularized B-spline transformation, in which the regularization strength was optimally chosen (Figure 2D).

**Figure 2:**
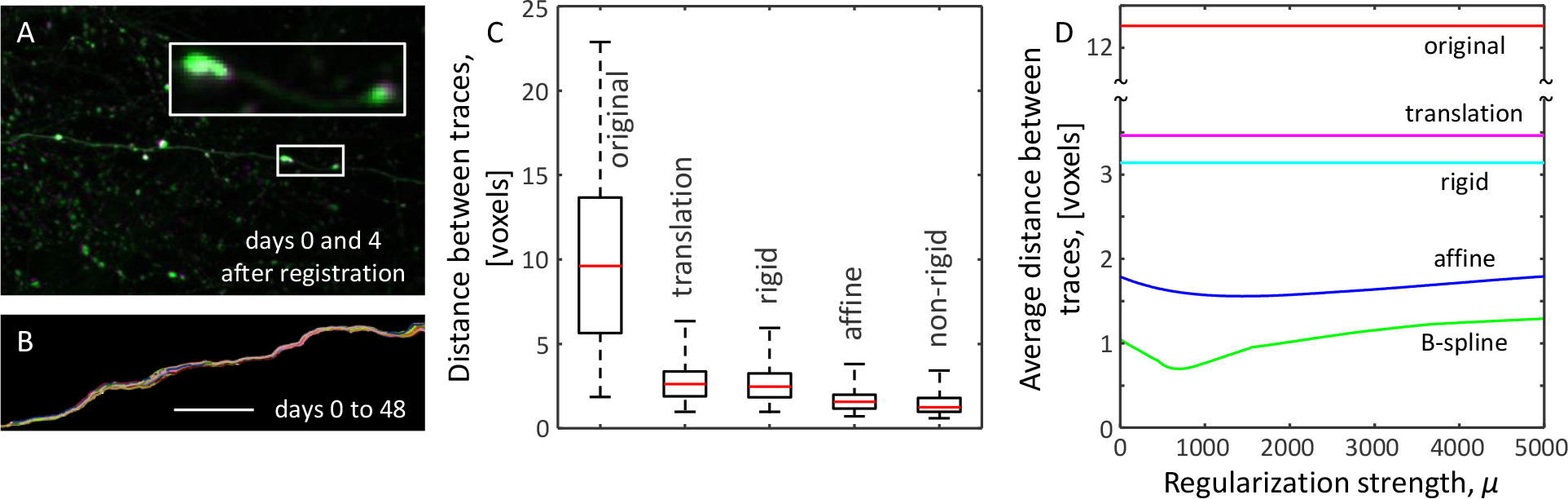
Validation of the registration procedure. **A.** After the registration (B-spline), the red-green overlap of the images from Figure 1 shows a significant improvement in alignment. **B.** The same trend is observed for the axon traces from Figure 1D. Scale bar is 25 μm. **C.** Box plots show the distances between traces of the same axons in subsequent time-lapse images before (original) and after the registration (translation, rigid, affine, and B-spline). **D.** Average distances between time-lapse traces for affine and B-spline transformations as functions of the regularization strength, *μ*. Corresponding distances in the original images and images registered with translation and rigid transformations are shown for comparison

### Registration enables accurate tracking of axonal boutons over time

In [6], boutons on the traced axons were automatically detected and tracked across imaging sessions in a semi-manual manner by using BoutonAnalyzer software (Figure 1A).

Using this dataset as a ground truth, we show that registration enables automated tracking of boutons (Figure 3B). To this end, we applied the Hungarian algorithm to the matrix of Euclidian distances between boutons in the original and registered images and compared the results to the ground truth. The results confirm that bouton tracking precision (0.71 ± 0.10, 0.97 ± 0.01, 0.98 ± 0.01, 0.994 ± 0.002, 0.997 ± 0.001) and recall (0.28 ± 0.06, 0.79 ± 0.05, 0.86 ± 0.04, 0.967 ± 0.012, 0.986 ± 0.005) gradually improves with increasing complexity of the transformations, and that boutons can be reliably tracked automatically following registration.

**Figure 3:**
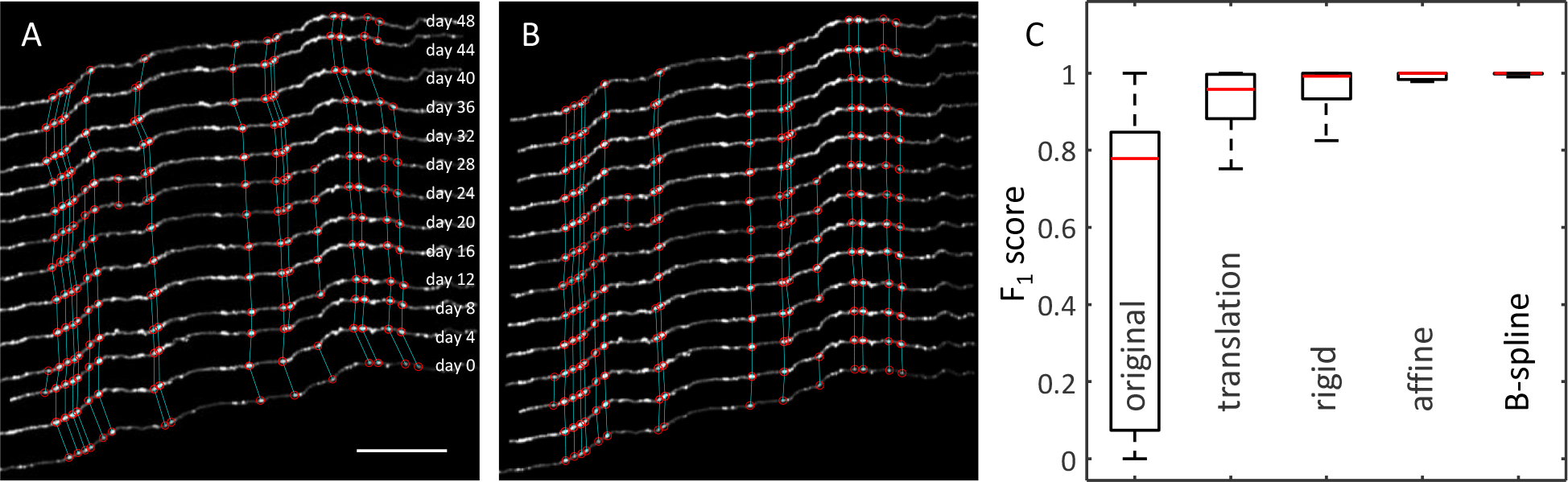
Registration enables accurate tracking of axonal boutons over time. **A.** An axon imaged in 13 sessions with a subset of boutons (red asterisks) detected and tracked (cyan lines) in a semi-manual manner with BoutonAnalyzer software. Axons are shifted vertically for better visualization. Scale bar is 25 μm. **B.** Images of the same axon after B-spline registration. Straightness of cyan lines is indicative of the registration accuracy. **C.** Registration makes it possible to track boutons automatically based on their proximity in the registered images. The *F*_1_ score is used to quantify the precision and recall of this automated bouton tracking procedure.

## CONCLUSION

We present accurate rigid and non-rigid methods for registering 3D optical microscopy stacks of images acquired in a time-lapse manner. The registration accuracy gradually improves with increasing complexity of the transformations, and sub-micrometer accuracy is achieved with the regularized B-spline (Figure 2). These results significantly improve upon the registration performed during image acquisition. As an example of utility of the proposed registration procedure, we applied it to track structural change in axonal boutons *in vivo*. Such changes are indicative of learning and memory formation in the brain, and accurate registration is essential to disambiguate true structural plasticity from differences due to image miss-alignment. We show that the presented method makes it possible to reliably track axonal boutons over the 48-day duration of the experiment with 99% precision and recall. This is a significant improvement over the 71% precision and 28% recall obtained based on the alignment done during the experiment. The proposed method is not specific to time-lapse images. It can be adapted to other essential medical imaging problems, including alignment of image planes within individual stacks and registration of overlapping stacks acquired in large-scale (e.g. whole brain) imaging experiments.

This work was supported by the NIH grant R01 NS091421.

